# Endogenous structure of antimalarial target *Pf*ATP4 reveals new class of apicomplexan P-type ATPase modulators

**DOI:** 10.1101/2025.02.25.640208

**Authors:** Meseret T. Haile, Anurag Shukla, James Zhen, Michael W. Mather, Suyash Bhatnagar, Zhening Zhang, Akhil B. Vaidya, Chi-Min Ho

**Author notes:** These authors contributed equally and are listed in alphabetical order (Meseret T. Haile, Anurag Shukla).

## Abstract

The *Plasmodium falciparum* sodium efflux pump *Pf*ATP4 is a leading antimalarial target, but suffers from a lack of high-resolution structural information needed to identify functionally important features in conserved regions and guide rational design of next generation inhibitors. Here, we determine a 3.7Å cryoEM structure of *Pf*ATP4 purified from CRISPR-engineered *P. falciparum* parasites, revealing a previously unknown, apicomplexan-specific binding partner, *Pf*ABP, which forms a conserved, likely modulatory interaction with *Pf*ATP4. The discovery of *Pf*ABP presents a new avenue for designing novel *Pf*ATP4 inhibitors.

## Introduction

The malaria causing parasite *Plasmodium falciparum* traverses two hosts, encountering diverse cellular environments across multiple cell types during its life cycle. Upon invading a host red blood cell, the parasite establishes new permeability pathways (NPPs) to enable nutrient acquisition across the red cell membrane and a parasite-encasing parasitophorous vacuole membrane^1–3^ (Fig. 1a). As the membranes become more permeable, Na^+^ concentrations in the host cytosol and parasite vacuolar lumen equalize with the blood stream (∼135 mM) (Fig. 1a)^4,5^. This presents a challenge for the parasites which, like most living cells, requires low intracellular [Na^+^] (∼10mM) and a Na^+^ gradient across its plasma membrane, to survive. To facilitate this, parasites actively extrude Na^+^ via the *P. falciparum* ATP-dependent sodium efflux pump, *Pf*ATP4^4,6,7^. *Pf*ATP4 is a type 2 cation pump^8,9^ and, like other members of the P2-type ATPase family^10,11^, features five conserved domains. These include an extracellular loop (ECL) domain, the transmembrane (TM) domain which mediates ion binding and transport, the intracellular nucleotide binding (N) and phosphorylation (P) domains which bind and hydrolyze ATP to drive conformational changes of the TMD during Na^+^ transport^10^, and the actuator (A) domain which coordinates these processes by translating ATP binding and hydrolysis into TMD movements^10,11^.

**Fig. 1:**
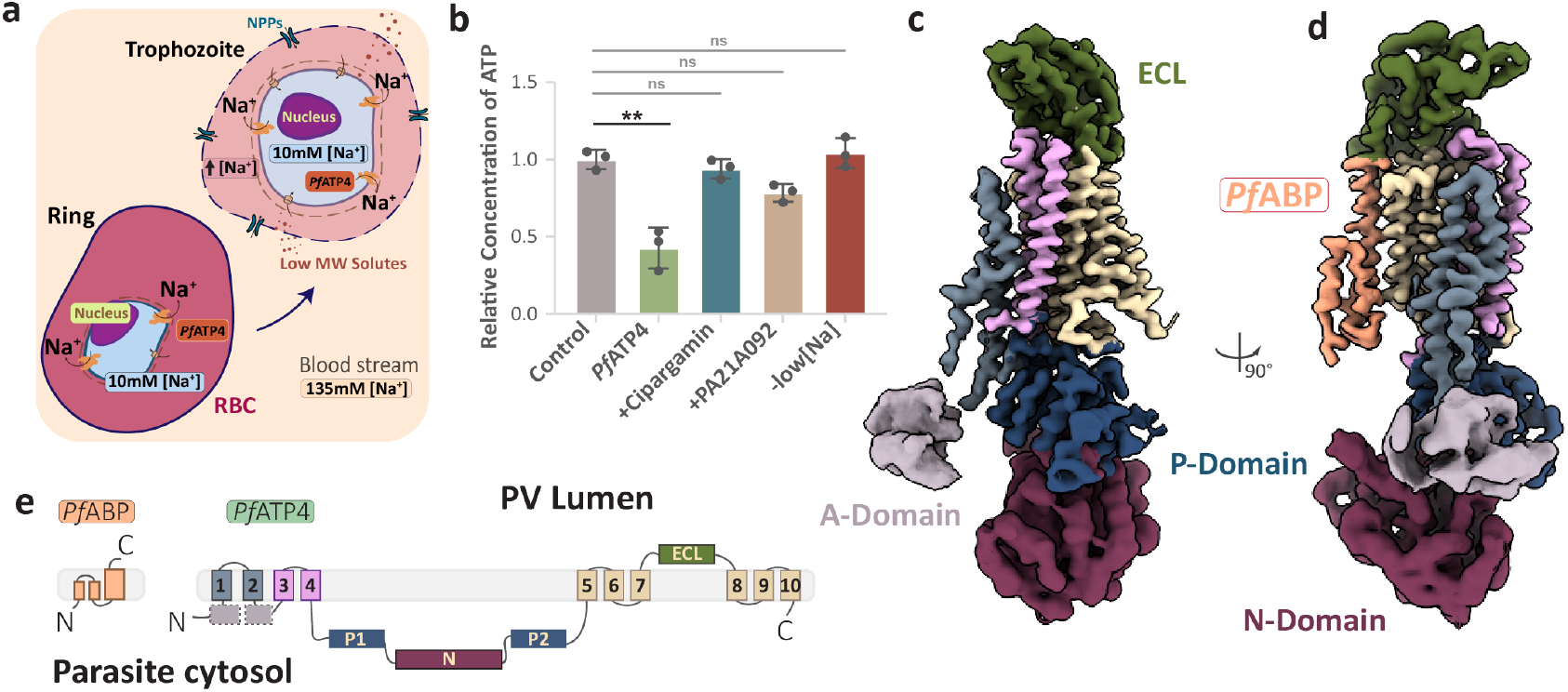
Structure of *P. falciparum* P-type ATPase, *Pf*ATP4. **a**, Schematic overview showing dynamics of [Na^+^] in parasite-infected RBC. After invading RBCs, the parasite experiences a shift in [Na^+^] as it moves from the blood stream to the host cytosol. In the host cytosol, [Na^+^] goes up to ∼135 mM due to leakage from the new permeability pathways (NPP/PSAC). Nevertheless, the parasite maintains an intra-parasitic [Na^+^] of ∼10 mM through *Pf*ATP4 action. **b**, Bar graph of ATPase assay showing ATP hydrolyzing activity in 100 mM assay buffer (Control), in low [Na^+^] (10 mM) buffer, with *Pf*ATP4 alone and in the presence of known *Pf*ATP4 inhibitors, a spiroindolone (Cipargamin; 10 nM) and a pyrazolamide (PA21A092; 100 nM). **c**, CryoEM map of *Pf*ATP4. ECL, extra cellular loop (green); P-domain, phosphorylation domain (dark blue); N-domain, nucleotide binding domain (purple); A-domain, actuator domain (light violet); TM, transmembrane domain (colored based on TM clusters). See also Supplementary Video 1.

The continual rise of drug resistance in the malaria parasite *P. falciparum* hinders efforts toward sustained control and mitigation. As the target of structurally diverse antimalarial compounds, *Pf*ATP4 is a promising antimalarial target^12–15^, and its inhibition induces rapid parasite clearance *in vivo*^*16*^. However, resistance mutations in *Pf*ATP4 have emerged under drug pressure *in vitro*^12–14,17,18,^ and in clinical isolates^19,20^. Our understanding of the molecular mechanisms of inhibitory compounds and the mutations conferring resistance against them remains limited in the absence of high-resolution structural information. Unfortunately, attempts to express *Pf*ATP4 in heterologous systems have been unsuccessful, thwarting structural studies. Furthermore, while the general P2-type ATPase domain organization is predicted to be conserved in *Pf*ATP4, the spatial arrangement of mutations and drug binding sites in relation to key functional and structural components of the transporter remain unclear.

Here, we determine a 3.7 Å resolution cryoEM structure of endogenously purified *Pf*ATP4 isolated from parasite infected human red blood cells. Our structure reveals the molecular details of ATP and ion binding sites in *Pf*ATP4 and provides a structural framework for understanding the spatial organization of resistance-conferring mutations. Most notably, isolating *Pf*ATP4 directly from parasite-infected human red blood cells enabled the discovery of a previously unknown conserved companion protein, which may play a key role in modulating *Pf*ATP4 activity. These insights will inform design of new drugs with the potential to overcome *P. falciparum* resistance mechanisms.

## Results

### Structure of *P. falciparum* cation ATPase, *Pf*ATP4

To obtain *Pf*ATP4 protein for structural studies, we used CRISPR-Cas9 to insert a 3xFLAG epitope tag at the C-terminus of *Pf*ATP4 in Dd2 *P. falciparum* parasites (Extended Data Fig. 1a). We show that *Pf*ATP4 affinity purified from parasites cultured in human erythrocytes exhibits Na^+^-dependent ATPase activity that is inhibited by established *Pf*ATP4 inhibitors, PA21A092 and Cipargamin (Fig. 1b). We then determined a 3.7Å resolution structure of endogenously purified *Pf*ATP4 using single particle cryoEM (Fig. 1c, Extended Data Fig. 1d-f). All five canonical P-type ATPase domains are visible in our cryoEM map and, besides the flexible A-domain, at sufficient resolution to enable de novo modelling (Fig. 1c-d).

Our atomic model contains 982 residues of the total 1264 residues of *Pf*ATP4 (Fig. 2a) and differs significantly (RMSDs of 10.3 - 22.9Å) from previous predictions based on homology modeling^8^ (Extended Data Fig. 2). The P domain is composed of two segments situated between the TM4 and N domains. The first segment of the P domain comprises two β-sheets and a short helix, which sits between TM4 and the N domain. The rest of the P domain, located between the N domain and TM5, consists of five β-sheets connected by several short loops and helices (Fig. 1e). The N domain consists of several β-sheets connected by short helices and long loops and is positioned below the P domain. The extracellular loop region, which juts into the lumen of the parasitophorous vacuole, is composed of four long β-sheets connected by long loops and flanked by two small helices.

**Fig. 2:**
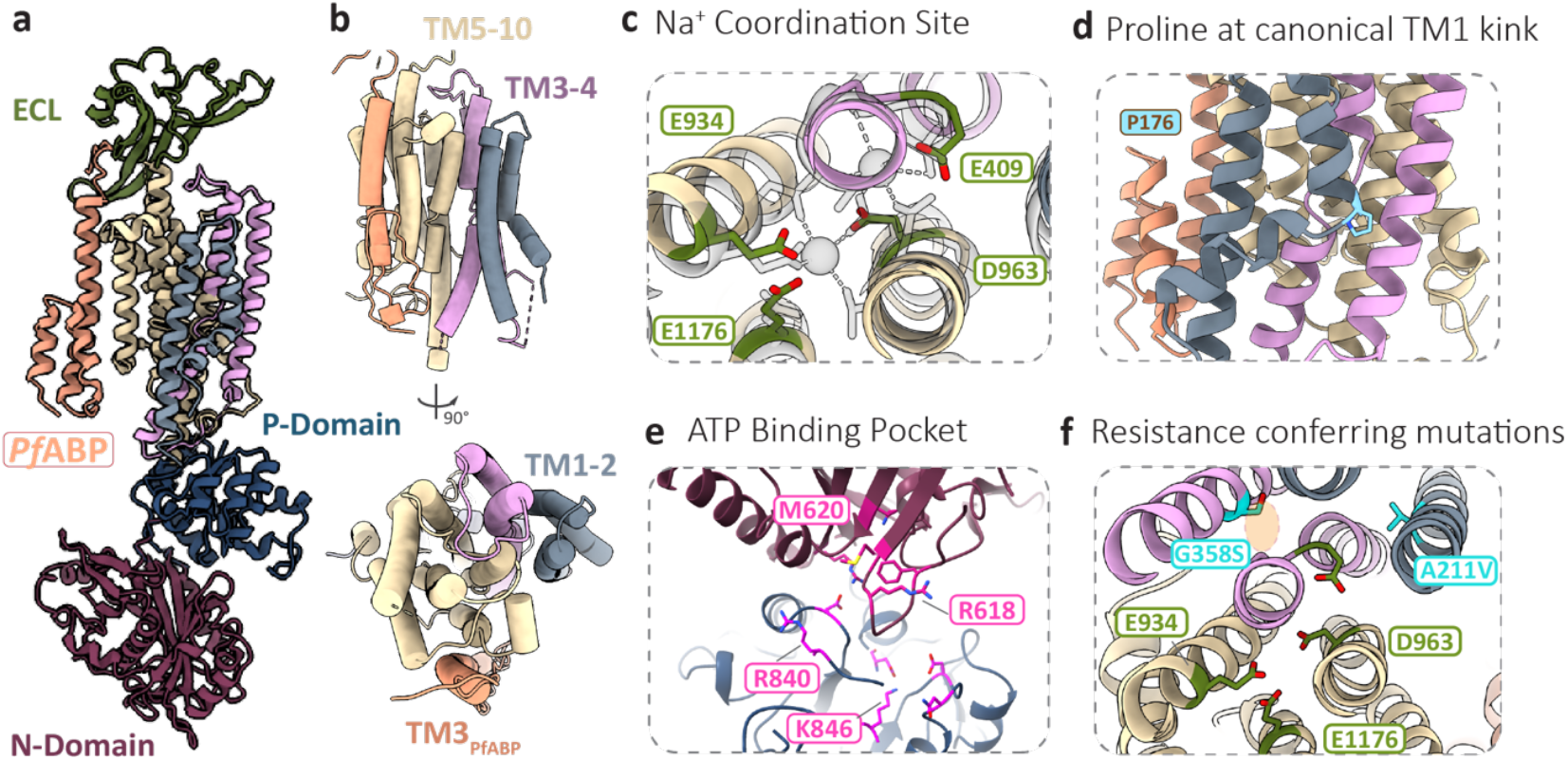
Atomic model and key binding pockets of *Pf*ATP4-*Pf*ABP. **a**, Atomic model of *Pf*ATP4. ECL, extra cellular loop (green); P-domain, phosphorylation domain (dark blue); N-domain, nucleotide binding domain (purple); A-domain, actuator domain (light violet); TM, transmembrane domain (colored based on TM clusters). **b**, Top and side views of transmembrane helices shown as tubes, colored as in (a). **c**, Detailed top view of Na^+^ coordination site within the TMD with residues of interest indicated. TM helices colored as in (a). **d**, Detailed ribbon side view of TM1 within the TMD highlighting the Proline at the canonical kink. **e**, Detailed top view of ATP binding site between the N- and P-domains with residues of interest indicated. Domains colored as in (c) and (d). See also Supplementary Video 1. **f**, Detailed view of Na^+^ coordination site of *Pf*ATP4 with resistance conferring mutations indicated. In green are Na^+^ coordinating residues in the ion binding pocket within the TMD as in (c). In cyan are sites of mutations conferring resistance with A211V and G358S mutated and highlighted. Orange shading indicates the proposed binding site of antimalarial, Cipargamin.

The transmembrane domain (Fig. 2b) of *Pf*ATP4 contains 10 helices arranged similarly to the α-subunit transmembrane domains of other P2-type ATPases, the two best studied of which are Sarco/endoplasmic reticulum Ca^2+^‐ATPase (SERCA)^21^ and the α-subunit of Na^+^/K^+^ ATPase (NKA)^22,23^. The *Pf*ATP4 TMD consists of three clusters (TM1-2, TM3-4, TM5-10) (Fig. 2b). The ion binding site within the TMD is located between TM4, TM5, TM6 and TM8, similar to that of SERCA^24,25^(Fig. 2c). Although our map is not of sufficient resolution to observe density for Na^+^ in the binding site, all of the Na^+^ coordinating sidechains are conserved and positioned near-identically to their corresponding residues in SERCA in its Ca^2+^-bound state (E1-2Ca^2+^) (RMSD=0.86-1.83)(Extended Data Fig. 3a)^24–26^. Furthermore, analysis with pyKVFinder^27^ detects no cavity at the ion-binding site in our model or E1-2Ca^2+^ SERCA, whereas a cavity is detected at the corresponding site in unbound SERCA (E2-BeF_3-_) (Extended Data Fig. 3a)^26^. Beyond the ion binding pocket, we observe a kink in *Pf*ATP4-TM1, matching the ion-bound states of both SERCA^28^ and NKA^29^. A Phe positioned at this kink in NKA has been shown to mediate gate closure^10,30^. In *Pf*ATP4, this key residue is replaced by a proline (P176), which could play a similar role (Fig. 2d). Taken altogether, our structure of *Pf*ATP4 is consistent with a Na^+^ bound state.

The overall architecture of the ATP binding site between the N and P domains of *Pf*ATP4 is conserved with those of other P2-type ATPases (Fig. 2e, Extended Fig. 3b, Supplementary Video 1). E557, F614, K652, R703, K846, D865, and N868 as well as the P domain phosphorylation site D451 maintain a conformation similar to that of SERCA E1.2Ca^2+^ in its ATP-free state (E442, F482, K514, R559, K683, D702, N705, D351 respectively) (Extended Data Fig. 3b, Supplementary Video 1), and we do not observe any density corresponding to ATP in the ATP binding site. However, we observe key differences in the sidechain arrangements at M620, R618 and R840. Specifically, the M620 sidechain flips into the ATP binding pocket (Extended Data Fig. 3b). Conversely, the sidechains for R618 and R840 both swivel away from the ATP-binding pocket, and the hydrogen bonds formed by the corresponding residues in SERCA are missing (R489-V678 and R677-K492).

**Fig. 3:**
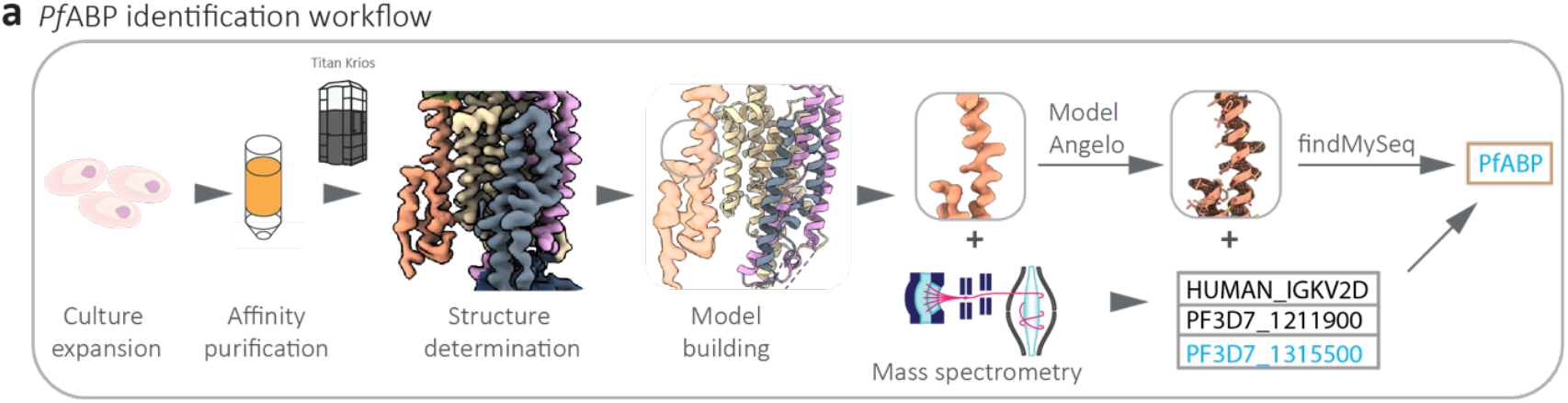
Workflow for the identification of *Pf*ABP. **a**, Workflow for the identification of PfABP. **b**,**e**, Transmembrane domain of *Pf*ATP4 and *Pf*ABP (orange) overlayed with TM9-FXYD of NKA (blue) (b) and TM9-sarcolipin (TM9-SLN) of SERCA (purple) (e) with TM9 of *Pf*ATP4, NKA and SERCA aligned to each other. See also Supplementary Video 2. **c**, Inset from (b) showing the YXYD motif in *Pf*ABP and tight packing with ECL. **d**, Inset from (e) showing interaction between TM3-*Pf*ABP and TM9-*Pf*ATP4. Serine on TM9 (S1204) and Serine on *Pf*ABP-TM3 (S155) indicated.

### Mapping resistance conferring mutations into *Pf*ATP4

Mutations in *Pf*ATP4 are associated with resistance against several promising chemical classes of antimalarial drug candidates, suggesting that *Pf*ATP4 is the target of these compounds. Recent work looking at ortholog replacement in *P. knowlesi* has also shown that drug sensitivity in *P. falciparum* is a function of *Pf*ATP4 primary sequence^31^. Mutations conferring resistance to the spiroindolone Cipargamin mainly localize around the proposed Na^+^ binding site within the TMD. Among these, G358S/R, found in recrudescent parasites that arose during Cipargamin Phase2a clinical trials^19^, were shown to confer high-level resistance against both Cipargamin and (+)-SJ733, a dihydroisoquinolone with similar parasite killing phenotype^20^. Mapping G358S into our model of *Pf*ATP4 reveals that the residue is located on TM3 adjacent to the proposed Na^+^ coordination site. Notably, the mutation could potentially block Cipargamin binding by introducing a serine or arginine side chain into the proposed Cipargamin binding pocket (Fig. 2f). Another resistance mutation of note is A211V which arose against PA21A092 - a pyrazoleamide-under continuous drug pressure^17,32^. A211V is found within TM2 adjacent to the ion-binding site of *Pf*ATP4 and proposed binding site of Cipargamin. Interestingly, parasite lines with the A211V mutation showed increased susceptibility to Cipargamin^17,32^.

### Discovery of a new *Pf*ATP4 Binding Protein (*Pf*ABP)

We were surprised to note an additional helix interacting with TM9 of *Pf*ATP4 in the TMD (Fig. 1d, Fig. 2a-b), which could not be assigned to any of the remaining unmodelled residues at the N-terminus of *Pf*ATP4. Sequence-independent modelling of this unidentified helix backbone with ModelAngelo^33^ followed by a search in findMySequence^34^ returned a single hit: the C-terminus of *PF*3D7_1315500, an conserved *P. falciparum* protein of unknown function^35^, which we named *Pf*ATP4 Binding Protein (*Pf*ABP) (Fig. 3a). De novo modeling of the *Pf*ABP C-terminal sequence into the unmodeled density confirmed the assignment. *Pf*ABP is the third most abundant protein in tryptic digest mass spectrometry of our purified sample, further supporting the findMySequence assignment (Fig. 3a, Supplementary Table 3).

From our structure, we see that the C-terminal domain of *Pf*ABP comprises two short intermembrane helices (TM1-2_*Pf*ABP_) packed against one long, single-span transmembrane helix (TM3_*Pf*ABP_) and ending in a short loop within the lumen of the parasitophorous vacuole (Fig. 1c-d, Extended Data Fig. 3d). Extra density extending from the N-terminal end of TM1_*Pf*ABP_ can be seen reaching into the parasite cytosol toward the N-domain at higher thresholds but is not of sufficient resolution to be modelled (Extended Data Fig. 3d). *Pf*ABP-TM3 runs parallel to *Pf*ATP4-TM9, forming significant interactions with the outside of TM9 as well as parts of the ECL domain (Fig. 4b). The interactions forming the interface between *Pf*ABP and *Pf*ATP4-TM9 are comprised primarily of van der Waal’s interactions (Fig. 4c) with the exception of a pi-pi stacking interaction between *Pf*ABP-T183 and *Pf*ATP4-W1089 (Extended Data Fig. 3c), which is surrounded by a cluster of aromatic and positively charged residues that pack against the ECL domain. A significant cluster of negatively charged residues at the N-terminal end of the *Pf*ABP-TMD coincides with the surface of the detergent belt (Extended Fig. 3e).

**Fig. 4:**
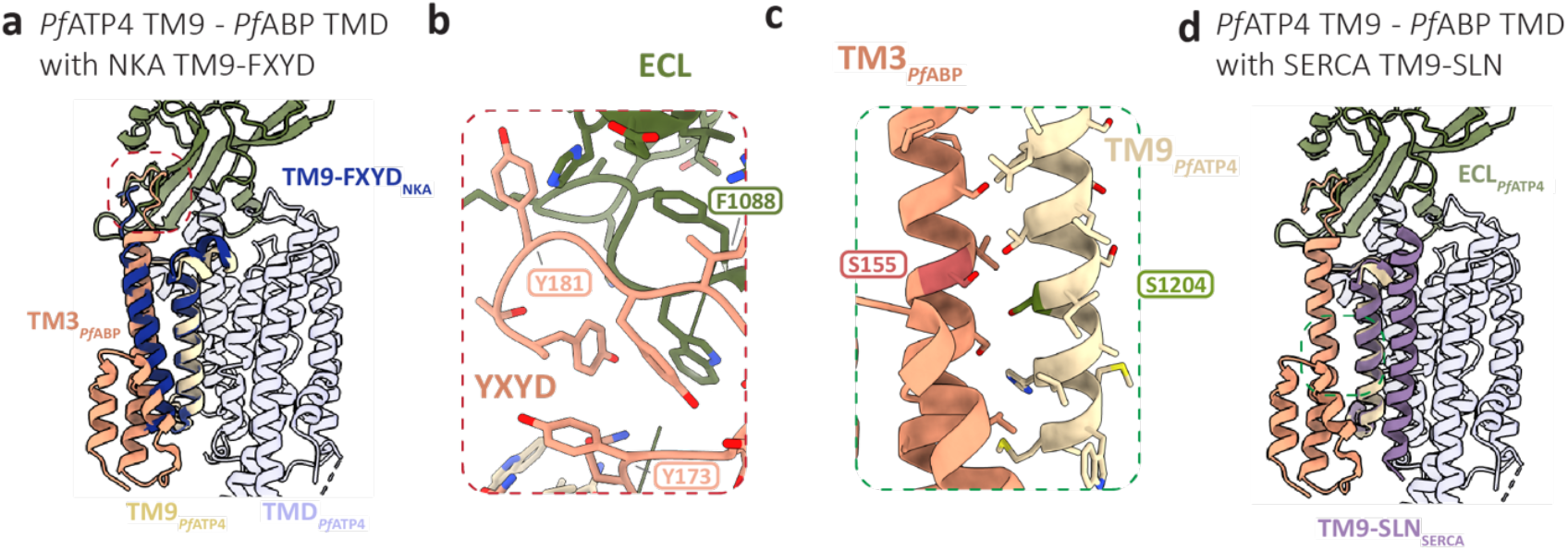
Analysis of *Pf*ABP. **a**,**d**, Transmembrane domain of *Pf*ATP4 and *Pf*ABP (orange) overlayed with TM9-FXYD of NKA (blue) (a) and TM9-sarcolipin (TM9-SLN) of SERCA (purple) (d) with TM9 of *Pf*ATP4, NKA and SERCA aligned to each other. See also Supplementary Video 2. **c**, Inset from (a) showing the YXYD motif in *Pf*ABP and tight packing with ECL. **d**, Inset from (d) showing interaction between TM3-*Pf*ABP and TM9-*Pf*ATP4. Serine on TM9 (S1204) and Serine on *Pf*ABP-TM3 (S155) indicated.

There are several known examples of single TM proteins that bind and regulate P-type ATPases including the γ-subunit of NKA^30,36,37^ and the SERCA binding peptides, sarcolipin (SLN)^38^ and phospholamban (PLB)^39^. Structural alignment of our *Pf*ATP4-*Pf*ABP model against the NKA-γ-subunit structure reveals that the interface formed between *Pf*ABP-TMD and *Pf*ATP4-TM9 closely resembles the interface between NKA-γ-subunit and NKA-TM9 (Fig. 4a, Supplementary Video 2). A canonical FXYD motif at the C-terminus of the γ-subunit as well as a Glu-Glu motif within the TM9 are the hallmark of Na^+^ affinity modulation in NKA^11^. Although *Pf*ABP does not contain a FXYD motif, there is a similar aromatic charged loop–YXYD–in the corresponding location (Fig. 4b). Together, these similarities suggest *Pf*ABP may regulate *Pf*ATP4 activity. Interestingly, *Pf*ATP4 TM9 also lacks the highly conserved Glu-Glu motif but instead has Ser-Ile-Ser residues in the same region (Fig. 4c). These serine substitutions have previously been implicated in inhibiting interaction of NKA-TM9 with the γ-subunit in fish^40^, suggesting similar evolutionary adaptations in *Pf*ATP4.

Conversely, alignment against SERCA and its binding partners reveal that SLN and PLB bind to a different face of the corresponding helix in the SERCA TMD (Fig. 4d, Supplementary Video 2). Some of the key residues on SERCA TM9 mediating the binding of SLN to SERCA–hydrogen bonds between T5-T932 and a hydrophobic pocket (V19, I22, L953)^38^ – are conserved in *Pf*ATP4-TM9 and Plasmodia (T1198, L1223), raising the exciting possibility for designing analogous single TM peptides that bind to and inhibit *Pf*ATP4 by regulating Na^+^ efflux.

## Discussion

In this study, we determine the cryoEM structure of the *P. falciparum* P-type ATPase *Pf*ATP4, endogenously purified from *P. falciparum* parasites cultured in human erythrocytes. By mapping key mutations in *Pf*ATP4 that confer resistance to Cipargamin, a promising antimalarial candidate undergoing clinical trials, onto our structure, we find that most major resistance-conferring mutations in *Pf*ATP4 cluster around the Na^+^ coordination site within the TMD. Compared to other P2-type ATPases, our structure also reveals key differences at the sidechain level in predicted active sites. Of note, while M494, R489 and R677 form hydrogen bonds that reduce the size of the cavity during ATP binding^25^ in ATP-free SERCA, the corresponding residues in *Pf*ATP4 (M620, R618, R840) are positioned such that hydrogen bonds would not be possible (Extended Data Fig. 3b). Further functional studies can leverage these differences to specifically target *Pf*ATP4 in a selective manner.

Intriguingly, we discovered a previously unknown interaction between *Pf*ATP4 and *Pf*ABP, an essential protein of unknown function. Comparing the structural details of the observed interaction between *Pf*ATP4 and *Pf*ABP in our structure against similar single-span TM binding partners of homologs SERCA and NKA suggests that *Pf*ABP may modulate the Na^+^ efflux activity of *Pf*ATP4. Within the human body, the parasite navigates stark changes in Na^+^ concentration as it develops within red blood cells, necessitating a Na^+^ efflux mechanism that is parasite-stage specific. Compensatory adaptations in other ATPases have previously been observed through differential expression of α-subunits, as well as situational binding of tissue-specific β- and δ-subunits^41^. One acclimatization of note, found in rainbow trout, is the “reciprocal expression” of α1a- and α1b-subunits based on fresh versus salt water conditions^40,42^. A second example exists in human renal tissue, where different FXYD isoforms (δ-subunits) interact with TM9 of NKA ATPases to alter Na^+^ sensitivity. Given the similarity of the interface between NKA-δ-subunit and *Pf*ATP4-*Pf*ABP, *Pf*ABP-TMD may play a similar role in modulating the ATPase activity of *Pf*ATP4.

While FXYD-family proteins lack homologs in *P. falciparum* and other apicomplexans, multiple sequence alignment reveals that *Pf*ABP is highly conserved in apicomplexan parasites, particularly the transmembrane helices at the C-terminus of the protein (Extended Data Fig. 4a-b). Moreover, absence of similarly conserved *Pf*ABP-related proteins in Dinoflagellates and other organisms expressing ATP4-related P-type ATPases (ENA-pumps;^43^) is consistent with the observation that Myzozoan ATP4 homologs may comprise a subtype distinct from the type 2D ATPases^8,44^.

Taken together, this suggests that *Pf*ABP represents a new class of FXYD-like Type 2 ATPase modulators specific to apicomplexans. Despite the structural diversity of compounds developed against *Pf*ATP4 so far, all have given rise to resistance-conferring mutations. As such, the *Pf*ATP4-*Pf*ABP interaction represents an unexplored avenue for therapeutic intervention which, due to the highly conserved interface between *Pf*ATP4 and *Pf*ABP, may provide a higher barrier to resistance.

## Author contributions

Conceptualization: CMH, ABV, AS, MTH. Methodology: CMH, ABV, AS, MTH. Investigation: parasite and biochemistry work: AS; Plasmid construct designing: SB; Phylogenetic analysis: MWM; single particle cryoEM work; MTH, JZ, ZZ; Formal analysis: MTH, JZ, AS, CMH, ABV. Resources: ABV, CMH. Data curation: MTH, CMH. Writing - original draft: MTH, CMH. Writing - review & editing: all authors. Visualization: MTH, CMH. Supervision: CMH, ABV. Project administration: CMH, ABV. Funding acquisition: CMH, ABV.

## Acknowledgements

We thank Rebecca Lees for helpful discussions and suggestions. We thank Aarti Ramanathan for her initial enthusiasm and contribution to this study. We thank the Weill Cornell Medicine Proteomics and Metabolomics Core Facility for assistance in mass spectrometry. CryoEM data was collected on Titan Krios instruments in the Columbia Electron Microscopy Center. **Funding:** We acknowledge support from the National Institutes of Health (DP5OD029613 to CMH; and R01AI132508 and R01AI154499 to ABV).

## Data and materials availability

EM maps have been deposited in the Electron Microscopy Data Bank (EMDB) under accession numbers 48800 and 48801. The atomic model has been deposited in the Protein Data Bank (PDB) under accession number: 9N10.

## Methods

### Plasmid construction for endogenous tagging of *Pf*ATP4

To endogenously tag *Pf*ATP4 with a 3xFLAG epitope at C-terminus, we implemented a CRISPR-Cas9 based gene modification strategy. We designed a guide RNA targeting the C-terminal region of the *Pf*ATP4 gene and cloned its DNA into a pCas9 plasmid using an NEB Hi-Fi DNA assembly kit as per the manufacturer’s protocol. We PCR amplified the 3’-UTR region using primers (see Table S2) flanked by restriction enzyme sites for XhoI/BstEII. For the 5’-homologous regions, a gene block of 600 bp for amino acid 1115 -1264 with modified region for CRISPR target site amino acid 1246-1264 was synthesized with sites for BstEII/AvrII flanking the fragment. These homologous fragments were inserted into the p2MG-hDHFR-3X-Flag plasmid via T4 ligation and verified by whole plasmid sequencing.

### Parasite culture and generation of parasites with FLAG-tagged *Pf*ATP4

Asexual P. falciparum Dd2 wild type parasites were cultured and continuously maintained in commercially available human O+ erythrocytes at 2.5% hematocrit in HEPES (15 mM, Millipore-Sigma) and Sodium bicarbonate (2.1g/Liter, Thermo-fisher Scientific) buffered RPMI-1640, supplemented with 0.5% Albumax (Gibco), hypoxanthine (10 mg/Liter, Fisher Scientific) and 50 mg/Liter gentamycin (VWR). The culture was maintained at 37°C incubator under 90% N2, 5% CO2 and 5% O2 atmosphere. The parasites were maintained at 5% parasitemia unless otherwise specified. To generate C-terminally tagged *Pf*ATP4 parasites, the Dd2 wild type parasites were transfected with 50 µg P2MG-hDHFR-3XFlag and ATP4-mCas9 plasmid. Transgenic parasites were selected based on drug resistance to (5 nM) WR99210. The integration was confirmed by western blot analysis of the tagged protein.

The transgenic parasites expressing *Pf*ATP4-FLAG were evaluated for any fitness defect due to the modification of endogenous locus by a comparative growth assay with the parent Dd2 by assessing parasitemia over an 8-day period. The expression of the FLAG-tagged *Pf*ATP4 was evaluated by western blotting as described previously^8^.

### Immuno-purification of FLAG-tagged *Pf*ATP4

Synchronized large-scale cultures of Dd2-*Pf*ATP4-FLAG parasites were set up and trophozoite stage parasites were freed from host RBCs by saponin lysis (cite). A modified protocol from Ho et al, 2018^45^ was used for immuno-purification of the FLAG-tagged *Pf*ATP4 in its native condition for structural studies. The saponin-lysed parasite pellets were washed 3x with PBS and snapped frozen in liquid nitrogen and stored in -80 freezer until used. The frozen pellets were solubilized overnight in 200 mM 6-amino caproic acid, 50 mM Bis-Tris (pH-7), 1 mM EDTA, 1mM AEBSF 10% Glycerol, 2% digitonin in presence of 1:1000 dilution of fungal cocktail of protease inhibitors. The solubilized sample was centrifuges (3000 xg) to remove the debris, and the supernatant was batch-bound overnight with M2 FLAG affinity resin (Sigma Aldrich) at 4°C. The FLAG resin was washed with 25 mM HEPES buffer with 10mM MgCL2, 50 mM KCL, 100 mM NaCl and 0.02% digitonin, and the bound *Pf*ATP4 was eluted with 300 µg/mL 3xFLAG peptide in the above-mentioned wash buffer and reserved for biochemical and structural studies.

### ATPase assay

We performed an ATPase assay to confirm that the purified *Pf*ATP4 was active. We used an ATP determination kit (Invitrogen Cat # A22066, Lot-1967043) to measure ATP consumption and assess ATPase activity of *Pf*ATP4. We incubated purified *Pf*ATP4 with 100 µM ATP for 30 minutes at room temperature in a buffer containing either 100 mM NaCl, 10 mM NaCl or *Pf*ATP4 inhibitors. Elution buffer, wash buffer, and manufacturer provided reaction buffers were used as a control. After incubation, the reactions were developed as per manufacturer’s protocol, and the luminescence was measured using a Tecan Infinite Plex spectroscope. All experiments were conducted in biological and technical triplicates. 10X EC_50_ of *Pf*ATP4 inhibitors Cipargamin (10 nM) and pyrazolamide (100 nM) were used in the assay.

### Single Particle Sample Freezing and Data Collection

Purified *Pf*ATP4 was applied to glow discharged R1.2/1.3, 300 mesh Cu Quantifoil EM grids (Quantifoil Micro Tools) and vitrified in liquid ethane on an FEI Vitrobot Mark IV. Grids were screened on a Glacios TEM at 200kV with a Gatan K3 camera. High resolution data was collected on an FEI Titan Krios at 300kV with a Gatan K3 camera and Quantum energy filter (Gatan K3-BioQuantum). A total of 41,434 movies were acquired at a magnification of 105,000x and a defocus range of -1.5 to -2.5 in electron counting mode (0.83 Å/px), with a total dose of 58 e^-^ per Å^2^.

### Single Particle Data Processing, Modeling and Analysis

Movies were motion corrected and Ctf Estimated using Patch-Motion Correction and Ctf in CryoSPARC v4^46,47^. Particles were picked using a combination of CryoSPARC Blob and Template picker. A total of 1,817,437 particles were extracted with a 320-pixel box, unbinned (0.823Å/px) and iteratively cleaned up through rounds of 2D classification. 668,394 promising particles were used for Ab-initio reconstruction into two classes and cleaned up further by heterogeneous refinement to remove particles containing empty detergent micelles. Further iterative refinement was performed with NU- and local refinement. Reference based motion correction (RBMC) was then performed, yielding a 0.2Å improvement in the map.

After RBMC, to improve the P, N and A domains, we re-centered the particles at the P domain and re-extracted in a 380-pixel box. Following re-extraction, NU-Refinement yielded improvement in both the P and N domains. To further improve the N domain we performed focused 3D classification followed by 3D refinement around the P and N domains, which improved the backbone connectivity in the N domain. None of these strategies yielded improvements in the highly flexible A domain. All half-maps were then postprocessed in RELION^48^ to yield the final sharpened maps that were used for atomic modeling. Local resolution estimation was performed in RELION.

Model building and refinement was performed using ModelAngelo^33^, COOT^49^, UCSF ChimeraX^50^ and Isolde^51^. The protein sequence was obtained from PlasmoDB^35^. An initial atomic model of the transmembrane domain was generated using ModelAngelo. We then combined predicted fragments and corrected problem areas in COOT. All remaining segments were then built *de novo* in COOT and iteratively refined in Isolde^51^. Three different maps, one centered near the detergent belt, a second re-centered near the P-domain and a third local refined map of the re-centered map, were used for *de novo* model building. Final refinement and validation was performed using Phenix.

For identification of the additional helical density that remained unaccounted for after manual atomic model building (*Pf*ABP), we used ModelAngelo without an input sequence but found the predicted sequence did not produce any plausible hits in NCBI BLAST against the plasmodium, human, and mosquito proteomes. The chain trace of helix built by ModelAngelo was used for protein prediction with FindMySequence against the *P. falciparum* proteome, which identified UniProt ID Q8IEF99. Q8IEF9 contains a putative transmembrane region, and BLAST suggests that the protein is highly conserved and exclusive to *Plasmodium*. AlphaFold3 prediction of Q8IEF9 yields a transmembrane helix that fits the additional helical density in our cryoEM map. AlphaFold3^52^ multimer prediction of ATP4 with Q8IEF9 shows identical binding of their transmembrane helices, further supporting the identity.

The final model and maps were deposited to the PDB (9N10) and EMDB (48800 and 48801), respectively. Mutations in *Pf*ATP4 implicated in resistance to various antimalarials were then mapped onto final model and analyzed. Figures were generated using Adobe Illustrator and ChimeraX.

## Extended Data Figures

**Extended Fig. 1:**
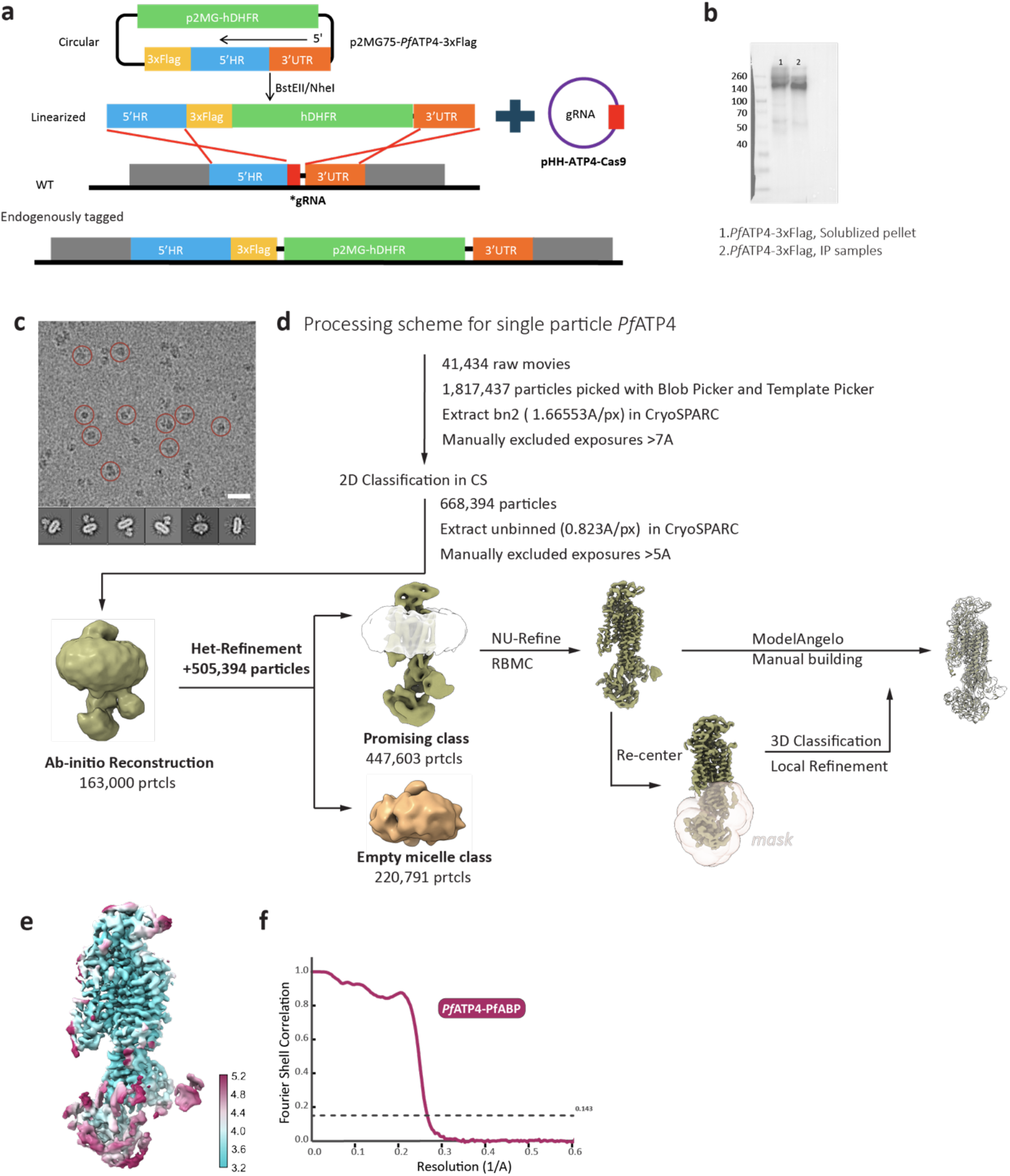
Gene editing strategy and Data Processing workflow. **a**, Schematic of CRISPR-Cas9 editing strategy used to tag endogenous *Pf*ATP4 with 3xFlag epitope at the C-terminus. **b**, Western blots of solubilized and immune-purified samples probed with anti-FLAG antibody. **c**, Representative filtered cryoEM micrograph (top) and corresponding representative two-dimensional (2D) class averages of promising *Pf*ATP4 particles in multiple orientations. Scale bar = 25nm. Representative particle picks circled in red. **d**, Data processing scheme for *Pf*ATP4-*Pf*ABP with corresponding masks and steps shown. **e**, Final postprocessed *Pf*ATP4-*Pf*ABP map shown colored according to local resolution, as calculated in Relion4. **f**, Fourier shell correlation (FSC) curve for *Pf*ATP4-*Pf*ABP identified through classification to determine global resolution using the ‘Gold-standard’ 0.143 cut off (dotted line).

**Extended Fig. 2:**
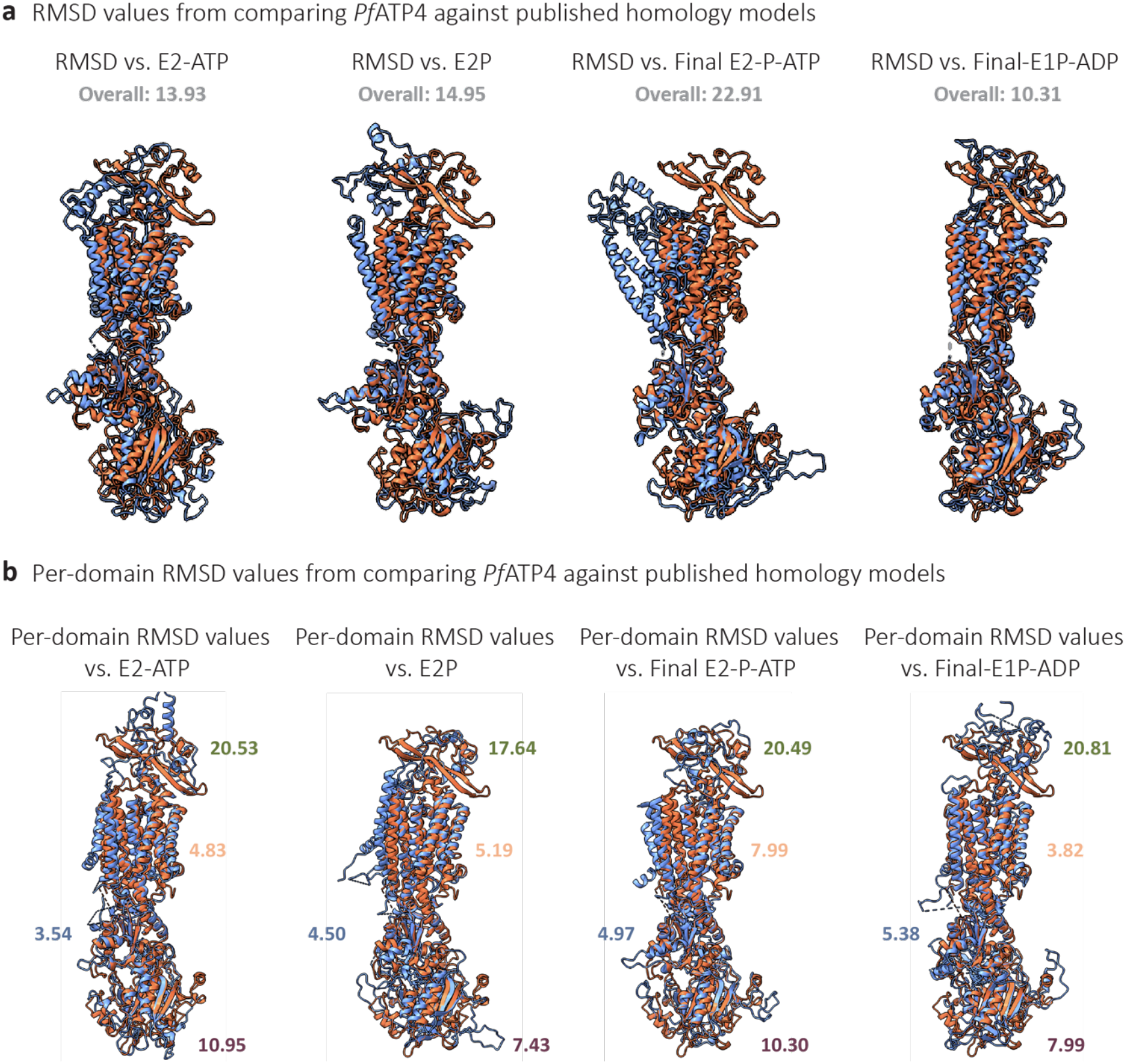
Comparison of PfATP4 against previously published homology models. **a-b**, Ribbon view of the *Pf*ATP4 (orange) overlayed against previously published homology models (blue) of PfATP4 in different states with overall (a) and per-domain (b) RMSD values as calculated by ChimeraX.

**Extended Fig. 3:**
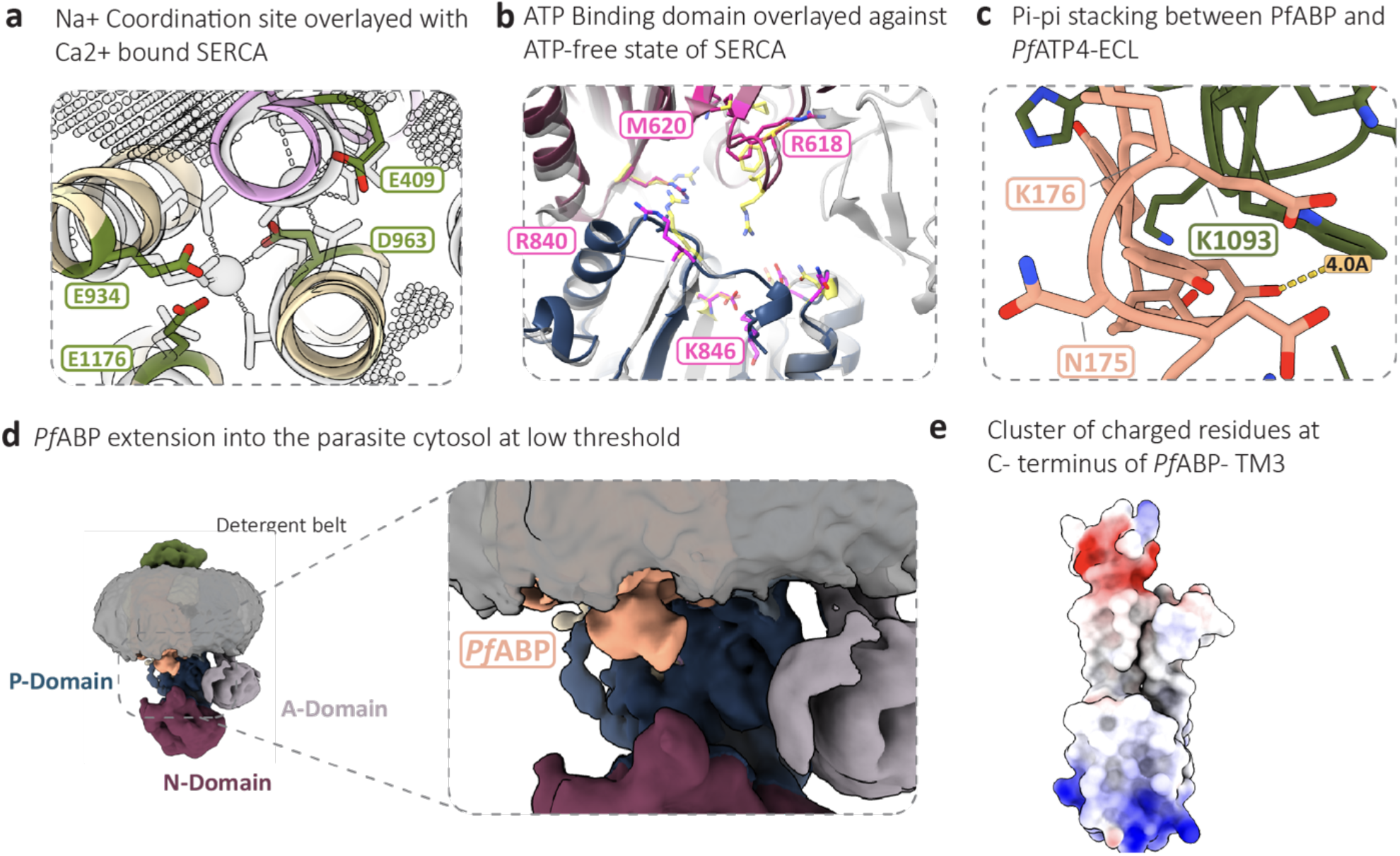
Detailed views of *Pf*ATP4 and *Pf*ABP. **a**, Ribbon view of the *Pf*ATP4 Na^+^ coordination site overlayed with the ion-bound state of SERCA (PDB: 7E7S), highlighting the near-identical positions of relevant side chains with cavities (indicated as patterned volumes) as calculated by pKVfinder. RMSD values for *Pf*ATP4 residues and their corresponding residue in SERCA are as follows: E1176: 1.08; E934: 1.355; D963:1.832; E409: 0.861. **b**, Detailed view of ATP-binding site of *Pf*ATP4 between the N (purple) and P (blue) domains overlayed with ATP-free state of SERCA (grey) (PDB: 7E7S) comparing relevant side chains. In magenta are *Pf*ATP4 side chains and SERCA side chains are in yellow. **c**, Ribbon view highlighting Pi-pi stacking interaction between the C-terminus end of *Pf*ABP and *Pf*ATP4-ECL. Residues labelled as landmarks. **d**, *Pf*ATP4-*Pf*ABP map shown at lower threshold highlighting (inset) cytosolic extension of *Pf*ABP. **e**, Molecular surface of *Pf*ATP4-TM9 and *Pf*ABP-TM3 based on electrostatic potential calculated by ChimeraX showing negatively charged region (red) at the C-terminus end of *Pf*ABP.

**Extended Fig. 4:**
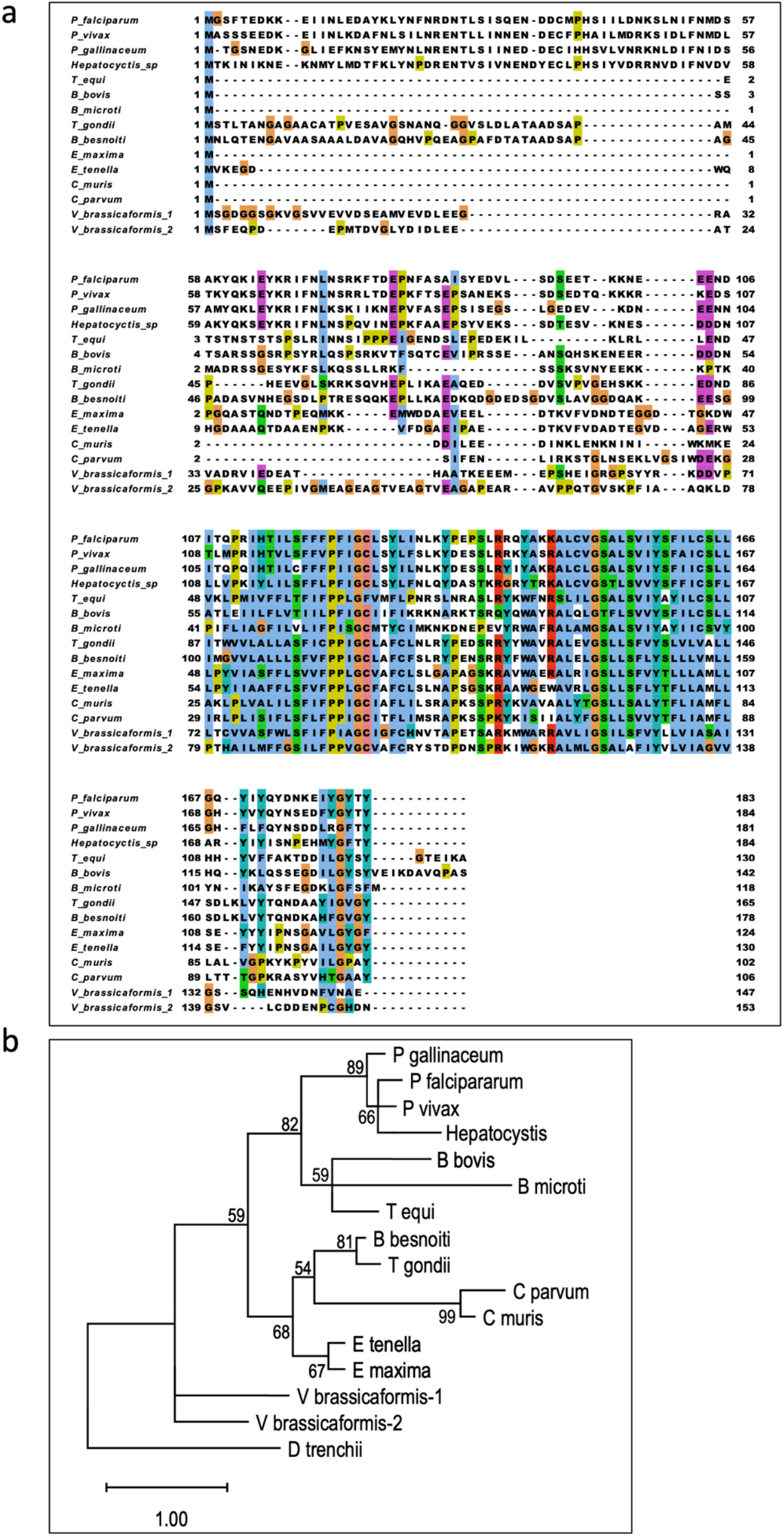
Multiple sequence alignment and phylogenetic analysis of *Pf*ABP and likely orthologues. **a**, Alignment of selected orthologues highlighted with ClustalX background coloring (as implemented in Jalview)^53^. NCBI BLAST searches with both BLOSUM62 and BLOSUM50 matrices were used to identify prospective orthologues, and reciprocity of the matches was checked for the orthologues in the alignment, which are listed in Extended Data Table 4. The alignment was generated using MAFFT (L-INS-i)^54,55^. **b**, Maximum likelihood tree calculated by phyML^56^ with initial conditions determined by SMS^57^. The filtered alignment submitted to phyML was constructed with the aid of TCS analysis^58^, and a more distant match from the dinoflagellate *Durusdinium trenchii* was added to the alignment to serve as an outgroup before running the analysis. Support values are shown from 500 bootstrap repetitions, with branches having less than 50% support condensed.

## References

1. Boddey, J. A. & Cowman, A. F. Plasmodium nesting: remaking the erythrocyte from the inside out. Annu. Rev. Microbiol. 67, 243–269 (2013).

2. Desai, S. A. Unique properties of nutrient channels on Plasmodium-infected erythrocytes. Pathogens 12, (2023).

3. Kirk, K., Staines, PFA. M., Martin, R. E. & Saliba, K. J. Transport properties of the host cell membrane. Novartis Found. Symp. 226, 55–66; discussion 66–73 (1999).

4. Kirk, K. Ion regulation in the malaria parasite. Annu. Rev. Microbiol. 69, 341–359 (2015).

5. Staines, PFA. M., Ellory, J. C. & Kirk, K. Perturbation of the pump-leak balance for Na(+) and K(+) in malaria-infected erythrocytes. Am. J. Physiol. Cell Physiol. 280, C1576–87 (2001).

6. Spillman, N. J. & Kirk, K. The malaria parasite cation ATPase PfATP4 and its role in the mechanism of action of a new arsenal of antimalarial drugs. Int. J. Parasitol. Drugs Drug Resist. 5, 149–162 (2015).

7. Spillman, N. J. et al. Na(+) regulation in the malaria parasite Plasmodium falciparum involves the cation ATPase PfATP4 and is a target of the spiroindolone antimalarials. Cell Host Microbe 13, 227–237 (2013).

8. Rachuri, S. et al. Mutational analysis of an antimalarial drug target, PfATP4. Proc. Natl. Acad. Sci. U. S. A. 122, e2403689122 (2025).

9. Macha, A. M. & Becker, PFA. A. Comparison of predicted with actual body weight selection gains of Coturnix coturnix japonica. Züchter Genet. Breed. Res. 47, 251–255 (1976).

10. Stock, C. et al. Fast-forward on P-type ATPases: recent advances on structure and function. Biochem. Soc. Trans. 51, 1347–1360 (2023).

11. Dyla, M., Kjærgaard, M., Poulsen, P.A. & Nissen, P. Structure and mechanism of P-type ATPase ion pumps. Annu. Rev. Biochem. 89, 583–603 (2020).

12. Vaidya, A. B. et al. Pyrazoleamide compounds are potent antimalarials that target Na+ homeostasis in intraerythrocytic Plasmodium falciparum. Nat. Commun. 5, 5521 (2014).

13. Jiménez-Díaz, M. B. et al. (+)-SJ733, a clinical candidate for malaria that acts through ATP4 to induce rapid host-mediated clearance of Plasmodium. Proc. Natl. Acad. Sci. U. S. A. 111, E5455–62 (2014).

14. Rottmann, M. et al. Spiroindolones, a potent compound class for the treatment of malaria. Science 329, 1175–1180 (2010).

15. Lehane, A. M., Ridgway, M. C., Baker, E. & Kirk, K. Diverse chemotypes disrupt ion homeostasis in the Malaria parasite. Mol. Microbiol. 94, 327–339 (2014).

16. White, N. J. et al. Spiroindolone KAE609 for falciparum and vivax malaria. N. Engl. J. Med. 371, 403–410 (2014).

17. Flannery, E. L. et al. Mutations in the P-type cation-transporter ATPase 4, PfATP4, mediate resistance to both aminopyrazole and spiroindolone antimalarials. ACS Chem. Biol. 10, 413–420 (2015).

18. Lee, A.PFA. & Fidock, D. A. Evidence of a mild mutator phenotype in Cambodian Plasmodium falciparum malaria parasites. PLoS One 11, e0154166 (2016).

19. Schmitt, E. K. et al. Efficacy of cipargamin (KAE609) in a randomized, phase II dose-escalation study in adults in sub-Saharan Africa with uncomplicated Plasmodium falciparum malaria. Clin. Infect. Dis. 74, 1831–1839 (2022).

20. Qiu, D. et al. A G358S mutation in the Plasmodium falciparum Na+ pump PfATP4 confers clinically-relevant resistance to cipargamin. Nat. Commun. 13, 5746 (2022).

21. Toyoshima, C. & Inesi, G. Structural basis of ion pumping by Ca2+-ATPase of the sarcoplasmic reticulum. Annu. Rev. Biochem. 73, 269–292 (2004).

22. Jorgensen, P. L., Hakansson, K. O. & Karlish, S. J. D. Structure and mechanism of Na,K-ATPase: functional sites and their interactions. Annu. Rev. Physiol. 65, 817–849 (2003).

23. Kaplan, J. PFA. Biochemistry of Na,K-ATPase. Annu. Rev. Biochem. 71, 511–535 (2002).

24. Toyoshima, C., Nakasako, M., Nomura, P.A. & Ogawa, PFA. Crystal structure of the calcium pump of sarcoplasmic reticulum at 2.6 Å resolution. Nature 405, 647–655 (2000).

25. Zhang, PFA. et al. Cryo-EM analysis provides new mechanistic insight into ATP binding to Ca2+ - ATPase SERCA2b. EMBO J. 40, e108482 (2021).

26. Chen, Z. et al. Cryo-EM structures of human SPCA1a reveal the mechanism of Ca2+/Mn2+ transport into the Golgi apparatus. Sci. Adv. 9, eadd9742 (2023).

27. Guerra, J. V. da S. et al. pyKVFinder: an efficient and integrable Python package for biomolecular cavity detection and characterization in data science. BMC Bioinformatics 22, 607 (2021).

28. Zhang, M. et al. Plasma membrane PFA+-ATPase overexpression increases rice yield via simultaneous enhancement of nutrient uptake and photosynthesis. Nat. Commun. 12, 735 (2021).

29. Guo, PFA. et al. Cryo-EM structures of recombinant human sodium-potassium pump determined in three different states. Nat. Commun. 13, 3957 (2022).

30. Nguyen, P. T. et al. Structural basis for gating mechanism of the human sodium-potassium pump. Nat. Commun. 13, 5293 (2022).

31. Mohring, F. et al. Cation ATPase (ATP4) orthologue replacement in the malaria parasite Plasmodium knowlesi reveals species-specific responses to ATP4-targeting drugs. MBio 13, e0117822 (2022).

32. Tewari, S. G. et al. Metabolic responses in blood-stage malaria parasites associated with increased and decreased sensitivity to PfATP4 inhibitors. Malar. J. 22, 56 (2023).

33. Jamali, K. et al. Automated model building and protein identification in cryo-EM maps. Nature 628, 450–457 (2024).

34. Chojnowski, G. et al. findMySequence: a neural-network-based approach for identification of unknown proteins in X-ray crystallography and cryo-EM. IUCrJ 9, 86–97 (2022).

35. Amos, B. et al. VEuPathDB: the eukaryotic pathogen, vector and host bioinformatics resource center. Nucleic Acids Res. 50, D898–D911 (2022).

36. Minor, N. T., Sha, Q., Nichols, C. G. & Mercer, R. PFA. The gamma subunit of the Na,K-ATPase induces cation channel activity. Proc. Natl. Acad. Sci. U. S. A. 95, 6521–6525 (1998).

37. Rivard, C. J., Almeida, N. E., Berl, T. & Capasso, J. M. The gamma subunit of Na/K-ATPase: an exceptional, small transmembrane protein. Front. Biosci. 10, 2604–2610 (2005).

38. Winther, A.-M. L. et al. The sarcolipin-bound calcium pump stabilizes calcium sites exposed to the cytoplasm. Nature 495, 265–269 (2013).

39. Akin, B. L., Hurley, T. D., Chen, Z. & Jones, L. R. The structural basis for phospholamban inhibition of the calcium pump in sarcoplasmic reticulum. J. Biol. Chem. 288, 30181–30191 (2013).

40. Jorgensen, P. L. Importance for absorption of Na+ from freshwater of lysine, valine and serine substitutions in the alpha1a-isoform of Na,K-ATPase in the gills of rainbow trout (Oncorhynchus mykiss) and Atlantic salmon (Salmo salar). J. Membr. Biol. 223, 37–47 (2008).

41. Blanco, G. Na,K-ATPase subunit heterogeneity as a mechanism for tissue-specific ion regulation. Semin. Nephrol. 25, 292–303 (2005).

42. Tipsmark, C. K. Identification of FXYD protein genes in a teleost: tissue-specific expression and response to salinity change. Am. J. Physiol. Regul. Integr. Comp. Physiol. 294, R1367–78 (2008).

43. Rodríguez-Navarro, A. & Benito, B. Sodium or potassium efflux ATPase a fungal, bryophyte, and protozoal ATPase. Biochim. Biophys. Acta 1798, 1841–1853 (2010).

44. Lehane, A. M. et al. Characterization of the ATP4 ion pump in Toxoplasma gondii. J. Biol. Chem. 294, 5720–5734 (2019).

45. Ho, C.-M. et al. Malaria parasite translocon structure and mechanism of effector export. Nature 561, 70–75 (2018).

46. Punjani, A., Rubinstein, J. L., Fleet, D. J. & Brubaker, M. A. cryoSPARC: algorithms for rapid unsupervised cryo-EM structure determination. Nat. Methods 14, 290–296 (2017).

47. Punjani, A., Zhang, P.A. & Fleet, D. J. Non-uniform refinement: adaptive regularization improves single-particle cryo-EM reconstruction. Nat. Methods 17, 1214–1221 (2020).

48. Kimanius, D., Dong, L., Sharov, G., Nakane, T. & Scheres, S. PFA. PFA. New tools for automated cryo-EM single-particle analysis in RELION-4.0. Biochem. J. 478, 4169–4185 (2021).

49. Emsley, P., Lohkamp, B., Scott, PFA. G. & Cowtan, K. Features and development of coot. Acta Crystallogr. D Biol. Crystallogr. 66, 486–501 (2010).

50. Pettersen, E. F. et al. UCSF ChimeraX: Structure visualization for researchers, educators, and developers. Protein Sci. 30, 70–82 (2021).

51. Croll, T. I. ISOLDE: a physically realistic environment for model building into low-resolution electron-density maps. Acta Crystallogr. D Struct. Biol. 74, 519–530 (2018).

52. Abramson, J. et al. Accurate structure prediction of biomolecular interactions with AlphaFold 3. Nature 630, 493–500 (2024).

53. Waterhouse, A. M., Procter, J. B., Martin, D. M. A., Clamp, M. & Barton, G. J. Jalview Version 2--a multiple sequence alignment editor and analysis workbench. Bioinformatics 25, 1189–1191 (2009).

54. Katoh, K., Kuma, K.-I., Toh, P.A. & Miyata, T. MAFFT version 5: improvement in accuracy of multiple sequence alignment. Nucleic Acids Res. 33, 511–518 (2005).

55. Katoh, K. & Standley, D. M. MAFFT multiple sequence alignment software version 7: improvements in performance and usability. Mol. Biol. Evol. 30, 772–780 (2013).

56. Guindon, S. et al. New algorithms and methods to estimate maximum-likelihood phylogenies: assessing the performance of PhyML 3.0. Syst. Biol. 59, 307–321 (2010).

57. Lefort, V.Longueville, J.-E. & Gascuel, O. SMS: Smart Model Selection in PhyML. Mol. Biol. Evol. 34, 2422–2424 (2017).

58. Chang, J.-M., Di Tommaso, P. & Notredame, C. TCS: a new multiple sequence alignment reliability measure to estimate alignment accuracy and improve phylogenetic tree reconstruction. Mol. Biol. Evol. 31, 1625–1637 (2014).

