## Supplementary Tables 1-4 for "Endogenous structure of antimalarial target *Pf*ATP4 reveals new class of apicomplexan P-type ATPase modulators"

**This file includes:**

Figure Legends for Supplementary Videos 1-2

Supplementary Tables 1-4

**Supplementary Video 1: Detailed view of *Pf*ATP4 ATP binding site.** ATP-binding site of *PfA*TP4 between the N (purple) and P (blue) domains overlayed with ATP-free state of SERCA (grey) (PDB: 7E7S) comparing relevant side chains. First showing PfATP4 alone, then SERCA alone, followed by the two structures overlayed. In magenta are *PfA*TP4 side chains and SERCA side chains are in yellow. See also Fig. 2e and Extended Data Fig. 3b.

**Supplementary Video 2: Detailed view *Pf*ABP.** *PfA*BP (orange) and transmembrane domain (light violet) and extracellular loop (green) of *PfA*TP4 shown then overlayed with TM9-FXYD of NKA (blue) and TM9-sarcolipin (TM9-SLN) of SERCA (purple). All three aligned based on TM9 of *PfA*TP4, SERCA and NKA. See also Fig. 3a,d.

**Supplementary Data Table 1: List of Oligonucleotides used for inserting epitope tag onto PATP4**

| **Primer** | **Sequence** |
|---|---|
| Guide RNA | CTTAGATGAAATAGTACCAA |
| 3' UTR-F (XhoI) | TGTACATGTGGTAAAAATTTTGCTTG (612 bp) |
| 3' UTR-R (BstEII/NheI) | TTAAATCACCACTACAACCATCTTGGTAACC |
| | GACGCGAGGAAAATTAGCATGCATC*GCTAGC*tata<br>att*GGTTACC*AAGATGGTTGTAGTGGTGATTTAAC<br>ACTTGATGAAAATGGATGGTGTAGGCCAAAGG<br>ATAATAAAACGTCTGATGGATACAATGATGAAC<br>TTGAGGGAATATTAAAAAAAGGTTTTGAAGATG<br>TCACGGCAAAAGGTTCAAAACGTGGTAGAACA<br>ATGGCATTTATATCAGCTGTTTGGTGTGAAATG<br>CTTAGAGCTTATACAGTAAGAAGTTGGGAACCT<br>TTCTATAAAGTATTTAATAGAAACATGTGGATG<br>CATTTAGCATGTAGTATATCTGCAACTTTAACA<br>TTTCTTTCAACATGTATACCTGGTATTACTTCTA<br>TTTTGAATACAACATGTTTGTTATGGTGGCAAT<br>ATTTATTAGCTATATTTTGGGCTCTTTTAAATTT<br>ATTTCTTGACGAGATTGTTCCTAAAGTaATtTATc |

| | |
|---|---|
| | GtAGgAAgTAcATGACaATaAAaAAc*cctagg*AATGG AGACTACAAGGACGATGACGATAAAggtGATTA TAAAGACGATGACGATAAAggaGATTATAAAGA TGACGATGACAAATAA***cgtacg***tcgagttatataatatatttatg |

**Supplementary Data Table 2: CryoEM Data Collection and Processing**

| | ***Pf*****A*****TP4** | ***Pf*****A*****TP4 recentered on the P-domain** |
|---|---|---|
| **Data Collection and Processing** | | |
| Magnification | x150,000 | x150,000 |
| Voltage (kV) | 300 | 300 |
| Electron exposure (e-Å$^2$) | 60 | 60 |
| Defocus range | -1.5 to -2.5 | -1.5 to -2.5 |
| Pixel size (Å) | 0.823 | 0.823 |
| Symmetry imposed | C1 | C1 |
| Initial particle images (no.) | 1,817,437 | 1,817,437 |
| Final particle images (no.) | 447,603 | 447,603 |
| Map resolution (Å) | 3.7 | 3.8 |
| FSC threshold | 0.143 | 0.143 |
| EMDB | 48801 | 48800 |

**Supplementary Data Table 3: Top twenty proteins detected with tryptic digest mass spectrometry of affinity purified *Pf*A TP4**

| **Protein Group** | **Protein Names** | **Peptides** | **Intensity** |
|---|---|---|---|
| A0A075B6S2; | KV229_HUMAN; KVD26_HUMAN; KVD29_HUMAN | 2 | 353644000 |
| PF3D7_1211900.1-p1 | non-SERCA-type Ca2+ -transporting P-ATPase | 83 | 181981000 |
| PF3D7_1315500.1-p1 | conserved protein, unknown function | 9 | 35395200 |
| PF3D7_1436900.1-p1 | histidine triad protein, putative | 8 | 20104900 |
| PF3D7_1456800.1-p1 | V-type H (+)-translocating pyrophosphatase, putative | 19 | 17879000 |
| PF3D7_0722200.1-p1 | rhoptry-associated leucine zipper-like protein 1 | 41 | 13848600 |
| PF3D7_1309300.1-p1 | U4/U6 small nuclear ribonucleoprotein PRP3, putative | 8 | 8760720 |
| PF3D7_1343900.1-p1 | U4/U6 small nuclear ribonucleoprotein PRP4, putative | 32 | 8373780 |
| PF3D7_1468900.1-p1 | zinc finger protein, putative | 48 | 7930770 |

| | | | |
|---|---|---|---|
| P81605 | DCD_HUMAN | 3 | 7871470 |
| PF3D7_0804800.1-p1 | peptidyl-prolyl cis-trans isomerase | 7 | 7405420 |
| PF3D7_0818900.1-p1 | heat shock protein 70 | 22 | 7167370 |
| PF3D7_1347200.1-p1 | nucleoside transporter 1 | 4 | 6074750 |
| PF3D7_0517000.1-p1 | 60S ribosomal protein L12, putative | 7 | 5942120 |
| P68871 | HBB_HUMAN | 9 | 5786940 |
| PF3D7_1462800.1-p1 | glyceraldehyde-3-phosphate dehydrogenase | 17 | 5740020 |
| PF3D7_1415400.1-p1 | Btz domain-containing protein, putative | 30 | 5714070 |
| PF3D7_0818200.1-p1 | 14-3-3 protein | 10 | 5617530 |
| P11142 | HSP7C_HUMAN | 7 | 5582920 |
| PF3D7_1237700.1-p1 | conserved protein, unknown function | 6 | 5406680 |

**Supplementary Data Table 3: Non-abbreviated Versions of *Pf*A*BP homologs used for Phylogenetic Analysis with corresponding accession IDs**

| **Species** | **Gene locus** | **Uniprot ID** |
|---|---|---|
| *Plasmodium falciparum* | PF3D7_1315500 | Q8IEF9 |
| *P. vivax* | PVP01_1416400 | A0A1G4H4J0 |
| *P. gallinaceum* | PGAL8A_00511800 | A0A1J1GZ83 |
| *Hepatocystis sp. ex Piliocolobus* | HEP_00232400 | A0A653H359 |
| *Theileria equi* | BEWA_018410 | L0AVF5 |
| *Babesia_bovis* | BBOV_III010310 | A7APV3 |
| *B. microti* | BMR1_01G02745 | A0A1N6LX20 |
| *Toxoplasma gondii* | TGME49_255410 | A0A125YKX2 |
| *Besnoitia besnoiti* | BESB_083850 | A0A2A9MB78 |
| *Eimeria maxima* | EMWEY_00019600 | U6MBK4 |
| *E. tenella* | ETH2_1113800 | (C-term of U6L4Y9 ) |
| *Cryptosporidium muris* | CMU_014170 | B6AEX4 |
| *C. parvum* | cgd7_4040 | Q5CY19 |
| *Vitrella brassicaformis (1)* | Vbra_8004 | A0A0G4EP91 |
| *V. brassicaformis (2)* | Vbra_4845 | A0A0G4EE69 |
| *Durusdinium trenchii* | CCMP2556_LOCUS32409 | (CAK9066014.1)* |

*no Uniprot entry; NCBI ID shown
